# Identification of genes important for response of *Pseudomonas aeruginosa* biofilms to ciprofloxacin exposure

**DOI:** 10.64898/2026.05.27.728104

**Authors:** Meng Wang, Emma Holden, Muhammad Yasir, Sarah Bastkowski, A. Keith Turner, Leanne Sims, Matthew Gilmour, Ian Charles, Mark Webber

## Abstract

*Pseudomonas aeruginosa* is an opportunistic pathogen that can cause severe infections in immunocompromised individuals, such as patients with cystic fibrosis where it commonly forms biofilms. Ciprofloxacin is used extensively to treat *P. aeruginosa* infections, but its effectiveness can be significantly reduced due to biofilm formation. Although many individual genes associated with biofilm formation or ciprofloxacin resistance have been characterised, the genetic basis of *P. aeruginosa* biofilm fitness related to antibiotic challenge remains incompletely understood. In this study we employed a whole genome screen to assay the impact of gene disruptions or altered gene expression on survival of *P. aeruginosa* biofilms exposed to different concentrations of ciprofloxacin. Genes impacting fitness in the biofilm context were identified by comparing the biofilm samples to planktonic samples harvested at 12h, 24h and 48h with and without ciprofloxacin.

Genes associated with c-di-GMP regulation and Gac/Rsm signalling were identified as primary regulators for biofilm formation in the presence and absence of ciprofloxacin. In addition, a group of genes involved in respiration, metabolism (especially polyamine metabolism), and various transporter and efflux systems were identified as important for biofilm fitness.

Ciprofloxacin specifically imposed a selective pressure on flagellar function and Psl production which were essential for survival in early biofilms. Moreover, transposon insertions within the CPA gene clusters (PA5448-PA5451 and PA5455-PA5456) and the salvage peptidoglycan recycling pathway showed reduced fitness in late biofilms at high concentration of ciprofloxacin, indicating that cell envelope integrity is beneficial for mature biofilms. This study identifies important determinants of survival for biofilms at different stages of maturity in the presence and absence of ciprofloxacin and implicates potential therapeutic targets for antibiofilm drug development.

## Introduction

*Pseudomonas aeruginosa* is a Gram-negative opportunistic pathogen and a leading cause of severe infections, particularly in patients with cystic fibrosis (CF), burn wounds, or immunocompromising conditions [1, 2]. Fluoroquinolones, particularly ciprofloxacin, have been extensively prescribed for treatment of *P. aeruginosa* associated infection due to their potent bactericidal activity and favourable tissue penetration [3]. However, therapeutic failure is common, occurring not only in infections caused by ciprofloxacin resistant isolates [4] but also in those involving strains reported as susceptible according to susceptibility testing [5, 6]. Apart from conventional intrinsic and acquired resistance mechanisms such as efflux pump overexpression and mutations in the ciprofloxacin target-encoding genes (*parCE* and *gyrAB*) [7], formation of a biofilm also represents a significant cause of ciprofloxacin failure. Biofilms are defined as structured communities where bacteria adhere to surfaces or form suspended aggregates encased within a self-produced matrix composed of polysaccharides, proteins, extracellular DNA (eDNA) and lipids [8]. *P. aeruginosa* is exceptionally proficient at forming biofilms and most of the chronic infections it causes are associated with biofilms [9, 10]. The complexed biofilm matrix can act as a physical barrier that slows, or limits penetration of some antibiotics, and cells within a biofilm are often metabolically dormant or in a persistent state. This results in antibiotic tolerance of the biofilm community which can often be 100 to 1000 times higher than that of planktonic counterparts [11, 12].

Antibiotic exposure has also been shown to enhance biofilm formation, especially at subinhibitory concentration of antibiotics [13, 14]. For example, exposure to ciprofloxacin has been shown to trigger the bacterial SOS response, leading to the release of eDNA that reinforces the matrix surrounding surviving cells within the biofilm [15, 16].

Over the last few decades, extensive work has identified numerous genes and regulatory networks involved in biofilm formation, maturation and dispersal in *P. aeruginosa*, including those controlling EPS production [8], cyclic-di-GMP signalling [17, 18], and surface appendages [19, 20]. However, the genetic determinants that specifically facilitate biofilm survival upon exposure to ciprofloxacin are less well understood.

In our previous work, a global genomics-based approach termed transposon directed insertion-site sequencing with expression (TraDIS-*Xpress*) has been used successfully to systematically investigate the genetic basis of biofilm formation in *Escherichia coli* [21] and *Salmonella Typhimurium* [22] as well as survival to various antibiotics [23, 24]. In this study, we employed the same technique to investigate the effect of deletion or altered expression of all genes within *P. aeruginosa* on biofilm adaptation under ciprofloxacin treatment. This identified common mechanisms important for biofilm formation in all conditions but also revealed genes specifically important for survival of biofilms when exposed to ciprofloxacin. These findings provide new insights into the genetic basis of biofilm maintenance under ciprofloxacin and highlighted potential therapeutic strategies against *P. aeruginosa* biofilm-associated infections.

## Methods

### Construction of transposon mutant library of *P. aeruginosa*

Five different transposons were created for mutagenesis in this study. All were mini-Tn5 derivatives coding for gentamicin resistance, and each included a different outward-transcribing promoter (from *dnaD, lpx02, rplJ, rplK* or *fabI*) to create a cocktail of mutants with promoters of different strengths. The transposon DNA fragments were prepared by PCR using Q5 high-fidelity DNA polymerase (New England Biolabs) then purified and concentrated using a DNA clean and concentrator kit (Zymo Research). Transposomes were prepared by mixing the transposon DNA with EZ-Tn5 hyperactive transposase according to manufacturer’s protocol (Epicentre) except that the transposon DNA concentration used was 100 – 200 ng/µL. *Pseudomonas aeruginosa* strain PAO1 was transformed by electroporation with transposomes using modifications of a previously published method [25]. To prepare electrocompetent cells, 400 mL Lennox broth (5 g/L NaCl) were inoculated with colonies from a Lennox agar plate, and the broth culture was incubated overnight at 37°C. Then, at room temperature, cells were harvested by centrifugation at 10000g and suspended in 0.3 M sucrose. This was repeated twice, then the final cell pellet was suspended in 2 mL 0.3 M sucrose. During one of the above wash steps, the cell suspension was left suspended in 0.3 M sucrose at room temperature for at least 1 h. For each transformation, 100 µL cell of suspension were mixed with 2 - 5 µL transposomes. Transformations were performed in 2 mm electrode gap electroporation cuvettes using an electroporator set to 2.5 kV, 25 µF and 200 Ω. Immediately after the electric pulse the cells suspension was rinsed from the cuvette with 1 mL S.O.C. medium (New England Biolabs) and incubated at 37°C with orbital shaking for 1.5 – 2 h. These cultures were then spread on Lennox agar supplemented with gentamicin at 30 µg/mL in 23 cm2 bioassay dishes and incubated at 37°C overnight. The approximate numbers of colonies obtained were determined, then harvested by scraping with ~7 mL Lennox broth poured onto the agar surface. Glycerol was then added to the harvested cell suspension to a concentration of about 15%, and 0.5 – 1 mL aliquots were stored at −70°C. For each of the five mini-Tn5 transposons, electro-transformations were performed until 200 thousand to 350 thousand colonies had been harvested, to give a total of just over 1.4 million colonies representing putative transformants.

### Minimum inhibitory concentration (MIC) determination

The wild type PAO1 strain used to generate the transposon mutant library was also used to determine ciprofloxacin susceptibility. The minimum inhibitory concentration (MIC) was determined using the broth dilution method. An overnight culture was diluted to a concentration of approximately 10^5^ CFU/mL in Miller Hinton (MH) medium. The culture was supplemented with various concentrations of ciprofloxacin and 150 μL of the bacterial suspension was dispensed to each well of the 96-well plate. Concentrations of ciprofloxacin ranged from 0 to 2 μg/mL and three replicates were included for each concentration. The cultures were incubated at 37 °C overnight before susceptibility was determined.

### Growth of biofilms

The biofilm and planktonic samples in this study were recovered from the same bacterial culture grown in 6-well plates. Briefly, the *P. aeruginosa* transposon mutant library was inoculated (5 μl) into 9 mL of Mueller Hinton (MH) broth to achieve a final concentration of ~5×10^5^CFU/mL. Sterile glass beads (~5mm diameter; Sigma-Aldrich) were added to each well (35 sterile per well) serving as substrates for biofilm formation. Plates were incubated at 37°C at 60 rpm in a humidified chamber. After 12h, 24h and 48h of incubation, biofilm samples and planktonic samples of three independent replicates were collected for DNA extraction. For each replicate, identical cultures in two wells were combined. Planktonic cells from 3 mL of culture were harvested by sedimentation, and pellets were washed by sterile PBS for DNA extraction. For biofilm cells, glass beads collected from two wells were dip-washed with PBS to remove loosely attached cells. Biofilm-associated cells attached to the beads were then resuspended in PBS by vertexing and pelleted by centrifugation at 10000x g for 1 min. Ciprofloxacin was added to the culture medium at the required concentration at the time of inoculation.

### Preparation of DNA fragments for TraDIS-Xpress

Genomic DNA (gDNA) of biofilm and planktonic samples was extracted using the Quick-DNA Fungal/Bacterial Microprep Kit and Miniprep Kit (Zymo research) respectively. The preparation of DNA fragments for nucleotide sequence generation using the known sequences in the end of the transposon as an anchor-point has been described previously [26]. Briefly, 100 ng DNA of each sample was fragmented and tagged using the MuSeek Library Preparation Kit (Thermo Scientific). Following a purification step with AMPure XP beads, the DNA fragments containing transposon insertion junctions were selectively enriched by an initial PCR using a pair of custom primers. The first primer was biotinylated and annealed to the transposon sequence and a customed i7 primer annealed to the tagged sequence generated by the Mu transposase. The resulting PCR products were sequentially purified using AMPure XP beads and then streptavidin beads, ensuring high specificity and purification of the target amplicon. A second round of PCR was subsequently performed using custom i5 and i7 primers to incorporate adapter sequences for high-throughput sequencing using the Illumina method. Fragment sizes were measured using a TapeStation (Aligent) and DNA sequenced on an Illumina NextSeq500 using the NextSeq 500/550 High Output Kit v2.5 with 75 cycles.

### Data analysis and bioinformatics

Nucleotide sequences generated from the sequencer were matched to the PAO1 (NC_002516.2) reference genome and analysed statistically using the QuaTradis pipeline (https://github.com/quadram-institute-bioscience/QuaTradis). Planktonic samples from each condition and each time point were compared against corresponding biofilm samples and a threshold of corrected p value of ≤0.05, log|FC| ≥ 1, and logCPM > 6 used to filter hits for significance. Visual inspection of insertion site mapping used the *Artemis* genome browser [27]. Annotated locus tags in the PAO1 annotated genome were correlated to gene names with help from the Pseudomonas Genome DB of the Cystic Fibrosis Foundation [28].

## Results

### Generation and validation of the high-density TraDIS-*Xpress* mutant library of *P. aeruginosa*

A high-density TraDIS-*Xpress* library of *P. aeruginosa* PAO1 was generated by random mutagenesis using five different mini-Tn5 transposon derivatives, each with a different outward-transcribing promoter to provide a range of expression levels. Following transformation, each of the five transposon variants yielded between 200,000 to 350,000 individual colonies, to give a total of over 1.4 million harvested putative mutant colonies. Genomic DNA extracted directly from these mutant collections was subjected to nucleotide sequencing using the TraDIS method which generated 8.8 million sequence reads that matched the PAO1 reference genome. This allowed the mapping of about 850 thousand unique transposon insertion sites, or one insertion site every 8 bp along the 6.26 MB genome, the mapping data confirmed coverage of mutants all across the genome (Figure 1A).

**Fig 1.**
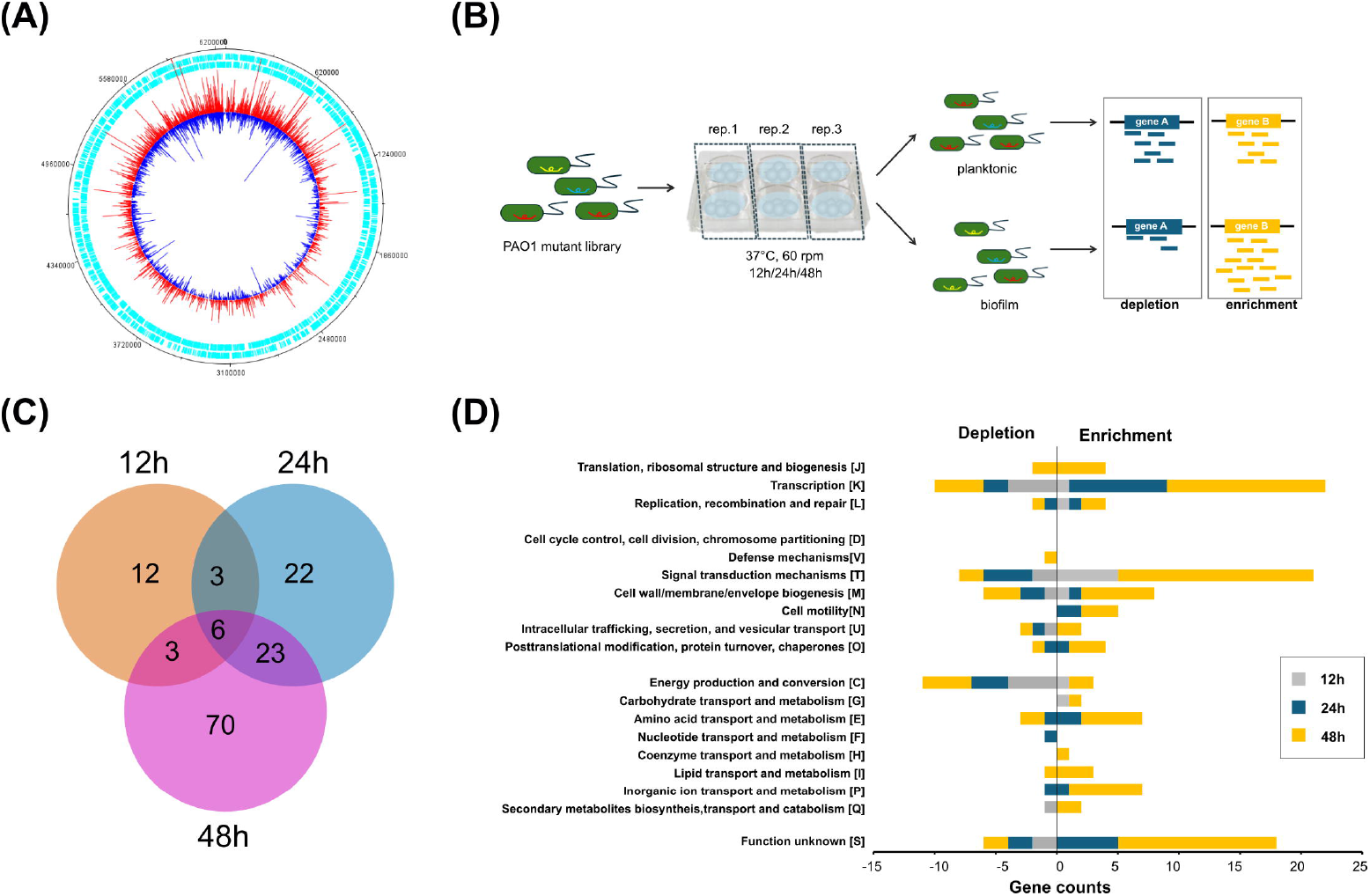

### The whole genome screen identified 139 genes that were involved in biofilm fitness

The *P. aeruginosa* transposon mutant library was cultured in nutrient broth with glass beads as biofilm substrates for 12h, 24h and 48h. Initially we sought to identify genes needed for biofilm formation in our system. At each timepoint, the planktonic culture and corresponding biofilm growing on the beads were collected and sequenced using the TraDIS-*Xpress* method (Figure 1B). Comparing the composition of transposon insertion mutants in the culture broth with that in the corresponding biofilm phase from the beads enabled the identification of genes involved in biofilm formation. Genes with significant depletion of insertion mutations in the biofilm samples (reduced fitness) were inferred to contribute to biofilm formation. Conversely, those with enriched insertions (increased fitness) in biofilm samples indicates that the disruption of these genes conferred a selective advantage under biofilm conditions.

A total of 139 genes were significantly differentiated between biofilm and planktonic culture at all three time points (Figure 1C; Supplementary dataset S1 & table S2) with mutants of 48 genes showing reduced fitness and 91 showing increased fitness in biofilm. Functional classification of these biofilm-associated genes was obtained using the Clusters of Orthologous Groups (COG) categories. Genes involved in signal transduction (T) and transcription (K) represented the largest functional groups, followed by a large group of genes associated with transport and metabolism of amino acids (E), inorganic ions (P), energy production (C) and cell envelope biogenesis (M). TraDIS-*Xpress* also identified many previously uncharacterised genes whose disruption resulted in either increased or reduced fitness in biofilm (Figure 1D), highlighting potential novel genetic determinants of biofilm development.

### Cyclic-di-GMP signalling is central to *P. aeruginosa* PAO1 biofilm formation

Cyclic-di-GMP signalling is a known regulatory pathway controlling the transition between planktonic motile growth and the sessile biofilm lifestyle in *P. aeruginosa*. About forty genes within the PAO1 genome have been discovered encoding active diguanylate cyclases (DGCs) for c-di-GMP synthesis and phosphodiesterase (PDEs) for degradation [29]. Consistent with the known role of c-di-GMP signalling in biofilm, TraDIS-Xpress analysis identified multiple c-di-GMP metabolic genes contributing to biofilm formation and revealed the dynamic involvement of these as the biofilms matured (Table S1). For example, genes in the *siaABCD* operon (PA0172–PA0169) were found to be involved at different time points of biofilm formation, with the signal sensor *siaA* identified as being important at 12h, the downstream kinase *siaB* identified at both 24h and 48h and adaptor gene *siaC* identified at 48h. Of these, insertion mutations within the *siaB* gene were enriched in the biofilms, while for the other *sia* genes, mutants were depleted. This fits with the known functions of the *sia* genes, where *siaB* encodes the protein inhibitor of the SiaD diguanyl cyclase leading to suppression of c-di-GMP synthesis and biofilm dispersal [30]. Transposon insertions within *rbdA*, whose products typically act as phosphodiesterase to degrade c-di-GMP, were also found to be enriched in biofilms at all three time points. We also identified significant enrichment of insertions within PA1850 at all three timepoints, these insertions were predominantly oriented in towards the immediately downstream PA1851 (Figure S1A). Since PA1851 was previously reported to be an active diguanylate cyclase (DGC) [31], the enrichment of insertions in PA1850 was interpreted to indicate that overexpression of PA1851 was beneficial for biofilm formation across the biofilm development.

In addition, transposon insertions within two other genes linked to c-di-GMP regulation were also identified at all three timepoints. They were *hsbA* (PA3347) and PA3623 (likely to disrupt neighbouring *rpoS*). HsbA is a part of the HptB signalling pathway which negatively regulates c-di-GMP production through the physical interaction between phosphorylated HsbA and diguanylate cyclase HsbD [32]. PA3623 is homologous to the lipoprotein NlpD/LppB of *E. coli* and no direct association between PA3623 and biofilm formation has been reported. However, the transcriptional regulator binding site of its downstream gene, *rpoS*, is located within the PA3623 coding region [33]. As *rpoS* encodes a global stress-response regulator and its mutant has been shown to elevate intracellular c-di-GMP and consequently promoted a hyper-biofilm phenotype in *P. aeruginosa* [34], the enrichment of insertions in PA3623 likely reflects disruption or altered regulation of *rpoS* rather than an effect of PA3623 itself.

The predominance of c-di-GMP related genes in the list of genes proposed to be important for biofilm formation confirms the central role of c-di-GMP signalling in regulating biofilm formation and validated the accuracy of the TraDIS-Xpress approach in identification of biofilm determinants.

### Microaerophilic respiration, matrix production and outer membrane homeostasis were important in early biofilms

In early biofilms, microaerophilic respiration was important for biofilm survival. Seven out of the fourteen genes identified at this stage were associated with microaerophilic respiration. These included genes encoding the high-affinity *ccb*_3_-2 cytochrome c oxidase isoforms (*ccoN2, ccoO2, ccoP2*), the biofilm-specific *cbb*_*3*_ subunit *ccoN4* (PA4133) together with its upstream regulator *mpaR* (PA4134) [35], as well as the global anaerobic regulator *anr* and oxygen-sensing component (*roxS*) of the RoxSR two-component system. Additionally, PA4131, encoding a putative iron-sulfur protein, was also required at 12 h, suggesting a potential role for redox homeostasis in early biofilm.

Whilst mutations in the *psl* operon were generally depleted in the biofilm, insertion mutants of *pslG* were enriched in the biofilm. The *psl* operon codes for Psl exopolysaccharide, and *pslG* encodes a periplasmic glycoside hydrolase that degrades Psl and regulates its accumulation. Thus, in our biofilm experiments, the PslG function for breakdown of the exopolysaccharide was detrimental to biofilm accumulation in the early stages. Proteins in the biofilm matrix represent another important structural component, mutants of secB (PA5128) and PA0943 were depleted in early biofilms suggesting protein export is crucial in early biofilms. SecB is a periplasmic chaperone of the Sec secretion pathway for translocation of unfold protein across inner membrane whereas PA0943 is responsible for correct localization of the XcpQ secretin in type II secretion system. Therefore, the identification of these two genes suggested that they contribute to biofilm fitness, likely through protein secretion to the biofilm matrix.

In addition, depleted insertions within *bamB* (PA3800) and PA4455 were identified in biofilms of 12h, indicating that outer membrane (OM) homeostasis was important for surviving in biofilm. BamB plays a role in the efficient assembly of the β-barrel outer membrane proteins, such as porins and fimbriae. PA4455 encodes a permease of ABC transporter MlaFEBD from *P. aeruginosa* which is responsible for phospholipid transport and contributes to the lipid homeostasis of the OM [36].

Generally, type III secretion system (T3SS) and biofilm formation are inversely regulated in *P. aeruginosa* representing a switch between acute and chronic (biofilm-associated) infection lifestyles. PA4595 encodes a putative ABC ATPase but is not genomically linked to a cognate membrane permease. Instead of a potential role as transporter, disruption of PA4595 has been reported to increase expression of type III secretion system (T3SS) independent of the known GacS/Rsm pathway [37]. Consistent with this, PA4595 mutants were depleted in biofilms in our TraDIS-Xpress data, suggesting that increased T3SS expression was not preferred to biofilm formation.

### There was a significant shift in the genetic requirements for biofilm at later stages

At later stages of biofilm development (from 24h to 48h), a group of hybrid sensor kinases and their signal transduction partners including *ladS*, PA1611, *retS, sagS, dspS and hptB* were identified to be important for biofilm. Amongst these, *lasS*, PA1611 and *retS* form a multi-sensor network to modulate the activity of GacS, subsequentially impacting *rsmY/Z* levels and shifting between motile and biofilm lifestyle. The orphan hybrid histidine kinase SagS also regulates biofilm post-transcriptionally through planktonic HptB-Gac/Rsm network or the surface-associated BifSR cascade where SagS activates the two-component system BifSR by direct interaction, which then drives the irreversible attachment and biofilm development [38]. DspS is a hybrid kinase sensor that sense the biofilm dispersal signal cis⍰2⍰decenoic acid (*cis*⍰DA) and mutation of *dspS* was found to coincide with larger microcolonies [39]. Consistent with this, transposon insertions within *dspS* were enriched in cells recovered from late-stage biofilms.

Various aspects of *Pseudomonas* metabolism are known to impact biofilm formation significantly. In this study, transposon insertions in multiple polyamine uptake (*spuG, spuH*) and catabolism genes (*pauA5, pauR* and *pauC*) were depleted in biofilms suggesting that polyamine metabolism was required at late stages of biofilm development. Supporting this observation, disrupted biofilm was previously reported in the polyamine-deficiency mutants of *Y. pestis*, whose polyamine pathway is similar as that of *P. aeruginosa* [40]. Mutants of *glpR* encoding a transcriptional repressor of glycerol uptake and metabolism were found to be enriched in biofilms, which in line with the previous report that activation of glycerol metabolism promoted biofilm formation [41].

Biofilm formation can provide oxygen-limited microenvironments which in *P. aeruginosa* can induce cyanide production. This was reflected by the involvement of different terminal oxidases and their regulators for respiration along with biofilm formation. At 12h, RoxS, a key two-component system regulating the cyanide-insensitive terminal oxidase *cioAB*, was required. Meanwhile, the identification of *mpaR* and *cooN4*, previously found to be induced by cyanide [42] also indicated production of cyanide. At 24h, CioA became essential for low-oxygen respiration along with the above terminal oxidases. In biofilms recovered after 48h, CioAB emerged as the sole terminal oxidase system required, indicating sustained accumulation of cyanide within the biofilm culture.

Several other known biofilm-associated genes responsible for biofilm matrix, metabolism and other regulators were identified at the late stages of biofilm formation with insertions in *amrZ, algZ, lapG, mexL, sutA, cmpX-crfX* and *bswR* enriched and insertions in *mexHI* depleted in biofilms. AmrZ acts as a transcriptional repressor of a DGC-encoding gene PA4843, and its disruption results in increased intercellular c-di-GMP levels, promoting biofilm formation [43]. LapG is a periplasmic protease cleaving large cell-surface adhesin CdrA that anchor cells to surfaces [44], consequently reducing stable biofilm attachment. The MexHI-OpmD transporter was required for biofilm survival to export the excessive reactive metabolite phenazine intermediate 5-methylphenazine-1-carboxylate (5-Me-PCA) during pyocyanin biosynthesis in biofilm [45]. Similarly, MexL has been recently characterised as a transcriptional activator of pyocyanin biosynthesis [46], and its disruption may therefore reduce production of 5-Me-PCA. CmpX-CrfX forms a small membrane⍰stress–responsive module and was likely regulate biofilm through PA1611 and GacS-Rsm network [47, 48]. BswR was previously found to regulate the switch between biofilm and T3SS through sRNA *rsmZ* [49]. SutA is a transcription factor that associates with RNA polymerase to maintain gene expression during slow growth and stress, previous study has implicated that mutation of sutA interferes with biofilm formation [50].

In contrast, mutants within a group of global regulators (*vfr, rpoS*) and the quorum sensing (QS) systems including *lasR, mvfR, pqsE* and *vqsM*, were significantly enriched in biofilms of 24h and 48h. This list expanded to include *pqsB, pqsD* and *rhlA* at 48h. Consistent with previous evolution studies where variants with point mutation in *lasR* and *rpoS* adapted to biofilm[34, 51], this finding may suggest that in the biofilm or chronic-like environment, selection favours mutants that shut down acute virulence and QS-coupled public goods to save energy and better tolerate stress.

### TraDIS-Xpress identified novel candidates that potentially impact biofilm fitness

Beyond previously characterised biofilm-associated genes, nearly half of the identified genes under selective pressure haven’t been previously directly associated with biofilm formation (Dataset S1 & supplementary table S1). Of these, insertions within PA0336, a conserved bacterial RNA pyrophosphohydrolase, were depleted in biofilms (Figure S2). Previous work has shown that PA0336 was a functional homologue of *E. coli* RppH and acted as a Nudix-type hydrolase involved in RNA turnover. Transcriptomics analysis indicated that PA0336 mutant affected expression of many biofilm regulatory gene such as the *rsmY, vfr, amrZ, rhlR* as well as pyocyanin production genes [52]. This suggested that perturbation in the nucleotide pool and gene expression may induce broader cellular changes that impact biofilm development. Mutants of the other genes identified in this study such as PA4673, PA4852, *mfd* and *rnd* that are involved in nucleotide modification, protein expression and translation may affect biofilm in a similar mechanism. Notably, insertion mutants of *mfd* encoding a transcription-repair coupling protein were enriched in biofilm across all three time points (Figure S2),unlike the other associated genes at either 24h or 48h, indicating the potential important role of *mfd* in biofilm fitness beyond its role involved in antibiotic susceptibility [53].

In addition, depletion of insertions in *oxyR* encoding an oxidative stress regulator and enrichment of insertions in *ppiD* encoding a periplasmic chaperone protein associated with envelope stress response [54] were identified in biofilms at late stages (figure S2), highlighting the role for stress adaptation in mature biofilms. In addition, several hypothetical genes were identified as being important for biofilm formation including PA0104, PA1728, PA1751, PA2229, PA2837, PA3225, PA4027, *argH*, PA5536(*dksA2*), PA5275. Although protein structures of ArgH and PA5536 have been solved, their exact functions are not very well known. PA2229 contains a molybdenum cofactor sulfurase C-terminal (MOSC) domain and MOSC domains generally act as sulfur-carriers for metal-sulfur cluster assembly. Therefore, PA2229 might be involved in some specialized metabolism or detoxification required in the biofilm environment. PA0104 locates upstream of the *coxBA-coIII* gene cluster, which codes for a terminal oxidase for energy production and is generally induced under starvation conditions and stationary phase growth. When investigating the insertion sites within PA0104, we found that the insertions were clustered at the 3’ end of gene PA0104, and showing a similar pattern with another cluster of insertions in the intergenic region between PA0104 and *coxB* (figure S2). Therefore, the disruption of PA0104 is likely to affect the expression of *coxBA-coIII* in late biofilm.

### Ciprofloxacin exposure selects for a distinct set of genes needed for survival in biofilms

Having established the baseline set of genes important for biofilm formation we sought to understand how ciprofloxacin impacted biofilms. To assess the biofilm-specific response to ciprofloxacin stress, ciprofloxacin was added to the starting culture at 0.5 x and 1 x MIC. Linear regression analysis to compare the different datasets revealed a pronounced shift in the genes under selective pressure within both biofilms and planktonic populations following ciprofloxacin treatment, particularly at high concentration (figure 2A & figure S3). To further investigate the specific genes under selective pressure when biofilms were exposed to ciprofloxacin we used the same workflow as above for ciprofloxacin-free conditions. The resulting biofilm-associated genetic determinants under ciprofloxacin exposure were then compared with those identified in the absence of ciprofloxacin (figure 2B). Unique genes identified under ciprofloxacin stress were investigated as candidate ciprofloxacin-specific genes relevant in biofilms. Consistent with the regression analysis, high ciprofloxacin stress imposed a strong selection on the biofilms and only a few genes identified by the 1xMIC exposure overlapped with those identified as important for survival in biofilms under ciprofloxacin-free conditions (Figure S3). Although transcription and signal transduction remained the dominant categories selected by ciprofloxacin treatment, functional enrichment analysis revealed ciprofloxacin reshaped the biofilm-associated genes with obvious changes in categories including cell motility (N) and cell envelope biogenesis (M), particularly at 1xMIC. (Figure 2C & Figure S3)

**Fig 2.**
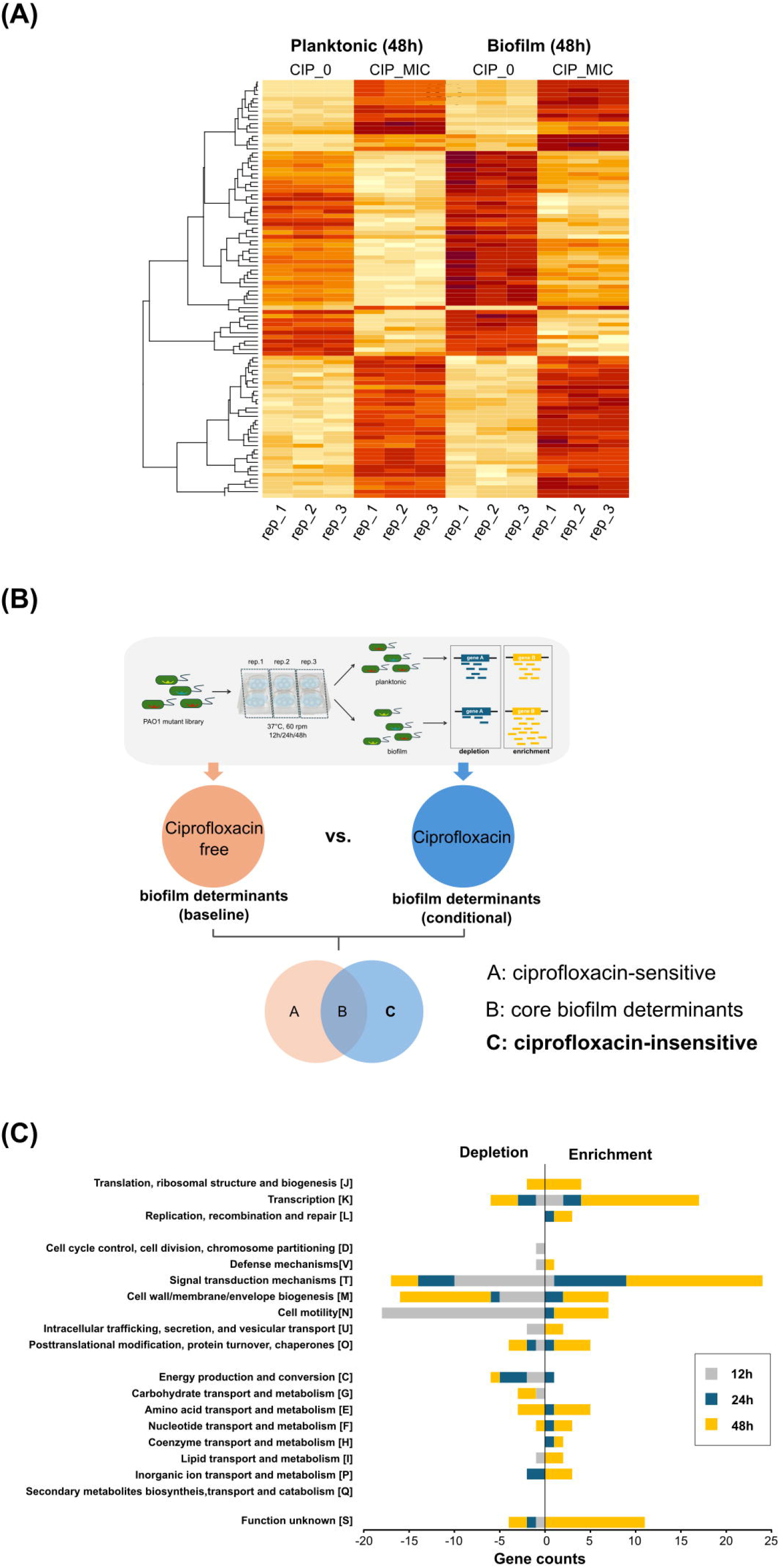

### Mutants of flagellar and exopolysaccharide Psl genes were negatively selected by ciprofloxacin in the early biofilms

Comparing with early biofilms under ciprofloxacin-free conditions, the most striking changes upon ciprofloxacin treatment was the selection of mutants within the major flagellar structural gene cluster and adjacent chemotaxis operon. At high ciprofloxacin concentration, insertions in 37 genes showed reduced fitness in the early biofilms, of which 22 were located within the flagellar and chemotaxis cluster (figure 2C & figure 3A), whereas only a few genes were identified in the absence of ciprofloxacin. The selection of mutations within flagellar genes in early biofilms was concentration dependent. (Figure 3A).

**Fig 3.**
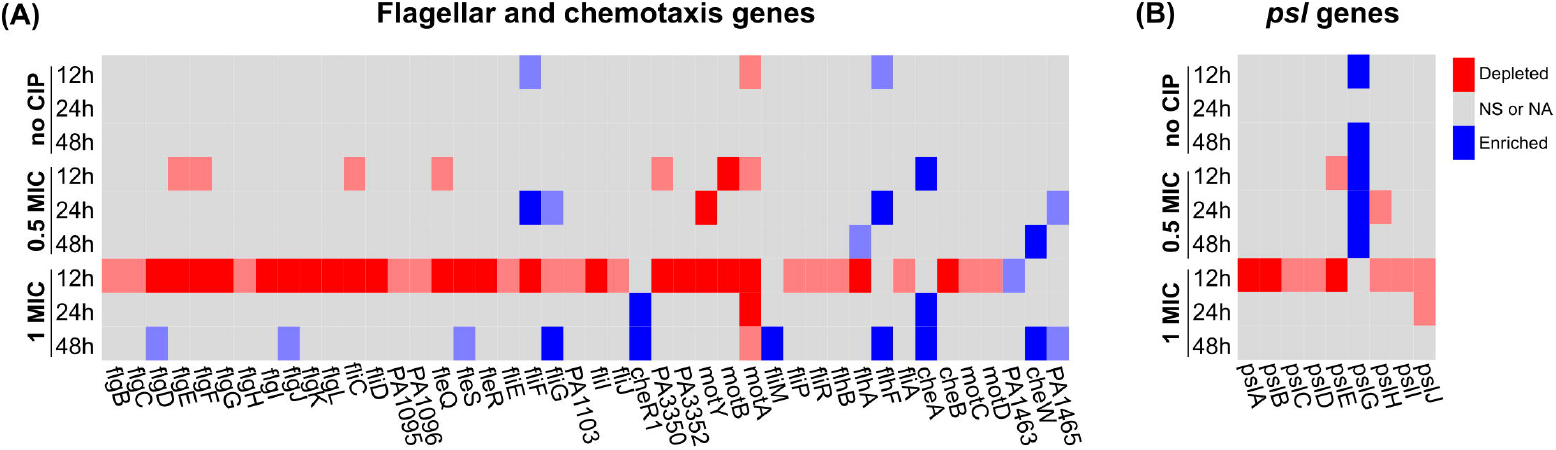

The core *psl* genes (PA2231-PA2240) within the Psl exopolysaccharide biosynthesis operon also displayed a strong ciprofloxacin-dependent selection (figure 3B). In the absence of ciprofloxacin none of the Psl biosynthesis genes were identified except for *plsG* where insertions were enriched in biofilms. However, as the concentration of ciprofloxacin increased, depletion of transposon insertions within genes including *pslE* and *pslH* at low concentrations and within a broader group of *psl* genes including *pslA, pslB, pslC, pslD, pslI* and *pslJ* at high ciprofloxacin concentrations were identified (Figure 3B).

### Biofilms also required multiple efflux system to mitigate ciprofloxacin toxicity

Multiple efflux systems including MexAB-OprM, MexXY-OprM and MexCD-OprJ were identified as contributing to ciprofloxacin tolerance both in biofilm and planktonic populations. Insertions leading to disruption of their negative regulators including *nfxB, mexR, mexZ, nalC, nalD*, and overexpression of PA5471 (including enriched insertions upstream of PA5471 and its leader peptide sequence) were dramatically enriched in both planktonic and biofilm samples upon ciprofloxacin treatment. Compared with the planktonic samples under ciprofloxacin stress, biofilm populations seemed to require higher expression of efflux systems as insertions within those efflux repressors were statistically more common than in planktonic samples. Consistently, insertions in the structural genes *mexA* and *mexB* were depleted in biofilms at 12h of half MIC and 1xMIC conditions respectively. This requirement in biofilms was both concentration and stage dependent. At low ciprofloxacin concentrations, only *mexZ* and *nalC* were identified at late stage of 48h while they were identified as early as 12h at high ciprofloxacin. In addition, disruption of *nalD, mexR, nfxB* and overexpression of PA5471 were all positively selected in 48h-biofilms at high ciprofloxacin.

### Mutants of several genes for flagellar, chemotaxis, and type IV pili were associated with ciprofloxacin tolerance in late-stage biofilms

TraDIS-Xpress analysis revealed mutants of additional genes including *cheR1, fliM, flhF, cheA, cheW* and PA1465 that didn’t appear significant in early biofilms as being enriched for mutants in late biofilms at the high ciprofloxacin concentration. Flagellar mutants are frequently reported to induce biofilm-like aggregation in response to environmental stress [55] and increased aggregation could enhance antibiotic tolerance of flagellar mutants. Furthermore, mutants of several flagellar genes (*flgD, flgJ, fleS, fliG*) that were initially depleted in early biofilms became enriched at 48h, though the insertion frequencies were relatively low (Figure 2A).

Beyond flagellar and chemotaxis genes, defects in type IV pili biosynthesis were also selected by ciprofloxacin in late biofilms. Although individual *fim* mutants were typically associated with impaired early biofilm formation, especially during initial attachment and microcolony development, three genes (*fimU, fimV* and *fimT*) showed enriched insertions in late biofilms in our TraDIS-data and their contributions to biofilm fitness varied depending on ciprofloxacin exposure. Disruption of *fimU* consistently benefited biofilm formation in dependent from ciprofloxacin stress. However, *fimT* mutants were no longer favoured in biofilms when exposed to high ciprofloxacin and *fimV* mutants were specifically selected by both low and high concentrations.

TraDIS-Xpress also identified PA5001 mutants as specifically enriched in biofilms of 12h and 48h under low ciprofloxacin stress, whereas no fitness difference was observed in the absence of ciprofloxacin. This indicates that the advantage conferred by PA5001 disruption is context-dependent and specific to low ciprofloxacin exposure. Given that loss of PA5001 has been reported to impair motility and alter biofilm architecture [56], its enrichment in biofilms may reflect adaptation to reduced ciprofloxacin penetration within the biofilm.

### LPS biosynthesis and the peptidoglycan salvage recycling pathway were essential for late-stage biofilm survival when exposed to the MIC of ciprofloxacin

While cell envelope integrity was a recurring theme for biofilm survival across our TraDIS-Xpress analysis (Figure 4A), a different set of genes associated with outer membrane and cell wall homeostasis were identified as essential for biofilm survival in the presence of ciprofloxacin. Those included *amgK, murU* and *anmK* from the salvage peptidoglycan (PG) recycling pathway [57] (Figure 4B), *mlaY* and *mlaZ* maintaining the membrane lipid asymmetry and *mepM* and *sltB1* involved in peptidoglycan biosynthesis and remodelling. Although PA5455 and PA5456 was identified under ciprofloxacin-free conditions, the requirement for A-band LPS biosynthesis intensified dramatically for surviving in late-stage biofilms especially at the higher ciprofloxacin concentration. Transposon insertions in a total of six genes from the eleven-gene LPS cluster (PA5455, PA5456, *wbpX, wbpY, wzt* and *wzm*) were significantly reduced compared to those in the corresponding planktonic cultures (Figure 4C).

**Fig 4.**
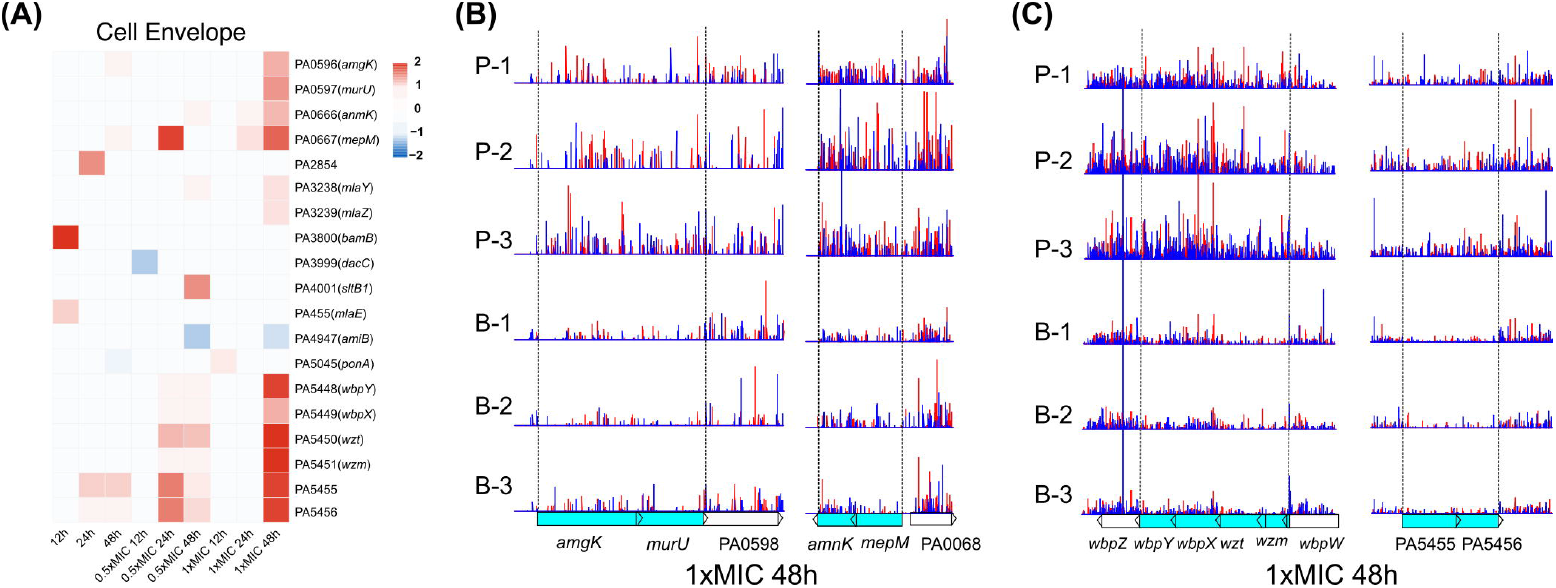

### Ciprofloxacin selected for altered functions in metabolism, transport, transcription, translation within biofilms

Fitness of several previously reported biofilm-associated genes were specifically identified as being important upon ciprofloxacin treatment. Insertions in *cafA* which is a member of the RNase G family linked to regulatory RNA (*rsmY/Z*) [58], were found to be depleted specifically in the 24-h biofilms under low ciprofloxacin condition and *rtcB* (PA4583) encoding an RNA ligase were found to be enriched in biofilms under ciprofloxacin conditions, which is in line with a previous study showing higher c-di-GMP level in the *rtcB* mutant [59]. In addition, mutants of *zapE* (PA4438), previously reported to reduce biofilm fitness against wild type PAO1, appeared to be enriched in biofilms at 24h under low concentration of ciprofloxacin. While c-di-GMP signalling pathway was still a central biofilm regulator in the presence of ciprofloxacin, the involvement of individual c-di-GMP metabolism genes were altered by ciprofloxacin exposure (Table S1). PA2771 encoding a DGC for c-di-GMP biosynthesis was no longer significant upon ciprofloxacin treatment, whereas PA5295 (*proE*) characterised as an active phosphodiesterase for c-di-GMP degradation [60] appeared important specifically under ciprofloxacin conditions with its mutants enriched in biofilm. Insertions in *sadC* (PA4332) which were enriched in late-stage biofilm under ciprofloxacin-free and low ciprofloxacin conditions were found depleted in early biofilms under high ciprofloxacin condition. Similar selection was also observed for the polyamine pathway in late biofilm. Instead of *spuG* and *spuH* which were identified without ciprofloxacin, the gene *spuF* for polyamine uptake were specifically required for biofilms at 48h at high ciprofloxacin.

Apart from genes where a previous link has been made to drug resistance, a range of genes were identified as having novel roles in survival of ciprofloxacin exposure in biofilms (Dataset S1, Table S3 & Table S4). Amongst these, insertions within *apaH*, PA3079, *greA* were significantly depleted in biofilm under both low and high ciprofloxacin conditions (Figure S4). ApaH hydrolyses the intercellular di-adenosine tetraphosphate (Ap4A), which accumulates under different types of stresses such as DNA damage [61-63]. GreA encodes a transcriptional elongation factor [64] that resumes the stalled RNA polymerase caused by environmental stress. PA3079 in *P. aeruginosa* sharing high similarity with the adhesin-associated MmpL efflux pump may be involved in exporting lipids that can contribute to surface attachment [65]. In contrast, mutants in several ribosome structural genes, including PA3179 and PA4932 (*rplI*), were enriched in biofilms (Figure S4), suggesting that altered translation may shift cells to adaptation in biofilms under ciprofloxacin treatment. Additional genes such as *rnk* and PA5130 whose mutants also adapted to biofilm under ciprofloxacin treatment (Figure S4). PA5130 is a rhodanese domain-containing protein potentially involved in sulphur metabolism and molybdenum cofactor biosynthesis, although its role in biofilm remains unclear.

## Discussion

In this study, we firstly used TraDIS-Xpress to investigate genome-wide gene fitness in the transposon library grown across multiple time points of biofilm development. To further investigate the impact of ciprofloxacin on biofilms, we then grew biofilms in the presence of ciprofloxacin and compared the identified gene determinants in the presence of ciprofloxacin with those in the absence of ciprofloxacin. Our analysis indicated that c-di-GMP regulation as well as the hybrid two-component system which controls the switch between acute and chronic infection through the Gac/Rsm network play central roles in biofilm formation and that these were conserved under both ciprofloxacin-free and ciprofloxacin stressed conditions. Flagellar function and Psl production were highly important for the early stages of biofilm formation and LPS biosynthesis was essential for late-stage biofilm survival at a high concentration of ciprofloxacin. Consistent with biofilm formation representing a lifestyle change, multiple genes associated with metabolism, transporter and quorum sensing showed reduced or increased fitness in biofilm. We also identified many hypothetical genes and previously uncharacterised genes as important for biofilm fitness, with or without antibiotic challenge (table S2-S4).

One key feature of TraDIS-*Xpress* distinct from other TnSeq approaches is that the outward-transcribing promoters fused with the transposon, enables identification of genes where expression changes contribute to fitness under given conditions. In our dataset, we observed biased insertions in PA1850, present within PA1848 and PA1849, being enriched in biofilm samples, which either represented disruption of PA1850 or overexpression of its downstream gene PA1851. To explore this, we created deletion mutants of PA1850 and PA1848 and examined their biofilm formation capacity by crystal violet assay. No significant difference of biofilm formation was observed between these mutants and wild type PAO1 (Figure S2). Together with the pervious characterisation that PA1851 encoded an active DGC [31, 66], we concluded that the activating insertions in PA1850-PA1848 act primarily by driving PA1851 overexpression. Notably, insertions orientated to the opposite direction were found exclusively within the intergenic region of PA1850 and PA1851 (figure S1A), leading us to propose that a potential transcriptional repressor of PA1851 might exist in *P. aeruginosa* and bind to this intergenic region. Further investigation is required to test this hypothesis.

The addition of ciprofloxacin to the biofilms led to a large change in the genes important for survival (figure S2C). Upon ciprofloxacin addition, nearly half of the genes identified under ciprofloxacin-free conditions (60 out of 139) were no longer important following antibiotic treatment. Insertions in a total of 163 genes and 148 genes were significantly differentiated between biofilm and planktonic samples at low ciprofloxacin and high ciprofloxacin, respectively, with 40 and 63 unique genes identified at each ciprofloxacin condition. The presence of ciprofloxacin not only impacted specific pathways important for biofilm but also demonstrated selectivity for specific genes within the same pathway, such as the selection to specific c-di-GMP metabolism genes, *fim* genes and polyamine genes under ciprofloxacin conditions. The most striking findings for biofilm important genes when stressed with ciprofloxacin were the identification of a group of functional genes related to Psl production and flagellar biosynthesis, and this requirement for biofilm formation seemed concentration dependent. Although the mechanisms for this selection remain unclear, it has been shown that deletion of Psl sensitized young biofilms to antibiotics [67] where lower concentrations of ciprofloxacin were required to eradicate existing biofilms formed by psl mutants than by wild type PAO1. It was also found that the protection of Psl to biofilm was effective for 24h-old biofilm against colistin but not for 48h and 72h of maturation, which was similar with our TraDIS data where psl genes were only essential at the early stage of biofilm formation under antibiotic treatment.

Flagella are well established as key determinants of early stage of biofilm formation in *P. aeruginosa*, particularly during initial surface attachment. However, in our dataset, flagellar genes were not prominently identified in biofilm at 12h under ciprofloxacin-free conditions. Instead, their importance was only identified in early biofilms in the presence of ciprofloxacin. One possible explanation for this discrepancy is the difference in biofilm growth rate. In the absence of ciprofloxacin, faster biofilm growth may mean that the 12-h time point no long captures the initial attachment phase, during which flagellar are critical. In contrast, addition of ciprofloxacin likely slowed the biofilm growth and extended the early-stage window, which allows the contribution of flagellar to be detected at 12h. Supporting this interpretation, genes associated with low-oxygen respiration such as *cioAB, roxSR* and *ccoN4* that were required by 24h-biofilms under ciprofloxacin exposure were already identified at 12h in ciprofloxacin-free condition, suggesting slower biofilms progress to oxygen-limited states in the presence of ciprofloxacin. To test whether this was true, we attempted to collect comparative samples at 6h but insufficient genomic DNA was obtained due to low cell density in the biofilm culture. Considering the gradient pattern observed for flagellar gene (and psl gene) fitness over time and along with the ciprofloxacin increase, this discrepancy was unlikely to be explained only by biofilm delay.

While the identification of classical biofilm associated genes supports the reliability of the dataset, we additionally selected eleven genes not previously implicated in biofilm formation to experimentally evaluate their functional relevance to biofilm (figure S4). In agreement with the TraDIS-Xpress data where insertions in PA3623 were enriched in biofilm, ΔPA3623 exhibited enhanced biofilm formation compared with wild type PAO1. Enriched insertions within an uncharacterized chemotaxis gene, PA1465, were also identified by TraDIS-Xpress. However, ΔPA1465 showed reduced biofilm formation; whilst this confirms a role for this gene understanding the mechanism behind this and how this is expressed in pure culture vs within a population requires more work. Not all the tested mutants where the TraDIS data suggested there was a fitness role for the gene in biofilm displayed a clear biofilm phenotype in the independently created mutants. This may reflect differences in the experimental set up where in the primary TraDIS screen mutants are within large pools and fitness benefits which can be dependent on phenotypes being supported by the neighbouring cells producing a product. Additionally, TraDIS-*Xpress* data are usually highly sensitive as results are based on competitive fitness changes of numerous independent mutants per gene in each condition and simpler biofilm assays with defined mutants can sometimes lack sensitivity to identify mild phenotypes.

In summary, a high-density PAO1 TraDIS-*Xpress* mutant library was generated for this study and identified many known biofilm determinants validating the sensitivity of our approach. Our data identified temporal signals for importance of various genes at different stages of biofilm maturity which provides new insights into the important events happening at these stages. In addition, we identified fundamental differences in the genetic landscape for genes needed for biofilms to survive in the presence of ciprofloxacin and again identified temporal and concentration-based specificities to the important genes. Collectively, this work illuminates more about the context specific genes and functions needed for *Pseudomonas aeruginosa* biofilms to survive ciprofloxacin treatment providing new information and novel potential targets which may have value for later exploitation.

## Data summary

Sequence data supporting the analysis in this study has been deposited in ArrayExpress under the accession number E-MTAB-16928. Results of the TraDIS-Xpress analysis are provided in Supplementary Dataset S1.

## Acknowledgements

We thank the Quadram Institute Bioscience Sequencing Core Facility for their contributions to this work.

